# Advancing AOPs for genotoxicity through quantifying KERs linking DNA alkylation to increase in mutations and chromosome aberrations

**DOI:** 10.64898/2026.11.11.765894

**Authors:** L.M.T. Winkelman, E. Demuynck, S.M. Bryce, M. Stępnik, A.I. Thienpont, F. Van Goethem, S. Majid, T.E. Pronk, M. Audebert, B. Mertens, J.B. Beltman

## Abstract

Quantitative Adverse Outcome Pathways (qAOPs) help translate mechanistic knowledge into quantitative toxicity predictions; however, their development is constrained by the limited availability and heterogeneity of quantitative data. Here, we investigated the feasibility of developing a qAOP for genotoxicity by modeling quantitative relationships linking DNA alkylation to increased mutations and structural chromosome aberrations through intermediate biological key events (KEs). We compiled and harmonized published concentration–response data from human lymphoblastoid TK6 cells exposed to ethyl methanesulfonate, methyl methanesulfonate, or N-ethyl-N-nitrosourea, covering eight genotoxicity assays representing five KEs with well-established dependencies. We evaluated sources of variability, compared points of departure (PoDs), and quantified relationships between successive events using response– response modeling. Substantial between-study variability could to a limited extent be explained through differences in experimental design while batch effects for individual studies and researchers were substantial. PoD estimates that were calculated from combined results of independent studies with similar experimental design, were often uncertain due to insufficient concentration–response characterization or conflicting results between studies. Nevertheless, response-response modeling suggested a threshold effect for certain key-event relationships (KERs). Specifically, DNA adducts seemed to be triggered at lower concentrations of alkylating agents compared to the subsequent KEs of DNA strand breaks and chromosome aberrations. Furthermore, these fitted response-response models can be used for prediction of downstream KEs from earlier ones. The general approach we developed for BMC analysis followed by response-response modeling is applicable to other AOPs as well. These findings highlight both the feasibility and limitations of using existing data for qAOP development.

## Introduction

Chemical genotoxicity assessment has largely relied on qualitative hazard-based approaches, in which the chemicals of interest are typically classified as either genotoxic or non-genotoxic (Menz et al., 2023; Omori & Hayashi, 2024). While effective for screening and classification, these rough classifications provide limited insight into genotoxic potency, dose/concentration–response behavior, and underlying mechanisms of action. Hence, quantitative understanding of how genotoxic effects evolve with concentration and how responses at different levels of biological organization relate to one another remains limited.

This limitation is particularly relevant because genotoxicity is inherently a dynamic biological process driven by sequential key events (KEs) that are interconnected (Cho et al., 2022; Demuynck et al., 2025). Depending on the mechanism of action, DNA lesions generally form rapidly following exposure to genotoxicants, whereas downstream processes such as DNA damage response activation, repair, mutation fixation, and chromosomal aberrations occur later across distinct temporal scales (Cho et al., 2022; Giles et al., 2025; Zentout et al., 2021). The magnitude and persistence of these events are further modulated by cellular defense mechanisms such as oxidative stress detoxification and DNA repair, DNA replication timing, and cell cycle regulation. As a result, measurements of genotoxic assays in isolation do not properly capture the mechanistic progression towards permanent DNA damage (Demuynck et al., 2025).

To better capture these interconnected and dynamic processes, we previously developed an AOP network for genotoxicity that integrates multiple genotoxicity-related AOPs reported in the AOP-Wiki and in the scientific literature into a comprehensible scheme (Demuynck et al., 2025). This AOP network connects several molecular initiating events (MIEs) to adverse outcomes (AOs) including increase in mutations and increase in structural chromosome aberrations (Cho et al., 2022; Paini et al., 2021). However, similarly to many toxicological endpoints, most AOPs for genotoxicity currently remain qualitative, limiting their predictive utility and regulatory applicability (Conolly et al., 2017; Perkins et al., 2019; Sinitsyn et al., 2022). Advancing from qualitative AOP descriptions towards quantitative AOPs (qAOPs) can improve the predictive utility of AOPs by characterizing the mathematical relationships among KEs (Perkins et al., 2019). When quantitative relationships are derived from evidence across multiple studies, however, differences in experimental conditions and biological systems may contribute to variability and uncertainty. This highlights the importance of evaluating comparability of data pertaining to the underlying KEs before integrating the evidence. For subsequent KER quantification, response–response modeling offers a promising and pragmatic strategy by focusing on associations between measured responses rather than administered dose / nominal concentration alone (Di Tillio et al., 2026; Doering et al., 2019, 2024).

Quantitative integration of available genotoxicity data presents two major challenges. First, genotoxicity studies have generally been designed to generate qualitative rather than quantitative data. As a result, concentration-response behavior is often insufficiently characterized, particularly for the low concentration ranges (low-response region) surrounding the expected point of departure (PoD). Second, there is substantial heterogeneity in experimental design across studies, which may strongly influence observed concentration–response relationships and inferred KERs. Differences in exposure duration, scoring methodology, time of harvest, cytotoxicity assessment, and assay configuration might all contribute to differences in measured genotoxic responses, thereby complicating cross-study integration and meta-analysis. Characterizing the impact of these experimental factors is therefore essential for identifying robust and transferable quantitative relationships between KEs.

Interest in qAOP development is growing (Spinu et al., 2020), alongside advances in mechanistic modelling of cellular DNA damage responses following chemical exposure (Burgers, Heldring, et al., 2025; Burgers, Sharma, et al., 2025; Heldring et al., 2022), which could be integrated into qAOPs. However, mechanistically integrated multi-endpoint datasets quantitatively linking early DNA damage to downstream genotoxic outcomes have, to our knowledge, not been established. Furthermore, no studies have systematically evaluated how experimental heterogeneity influences quantitative relationships between successive KEs within genotoxicity. Addressing these limitations is essential for developing robust, predictive and mechanistically interpretable qAOPs for genotoxicity assessment.

Within our previously developed AOP network for genotoxicity (Demuynck et al., 2025), the subnetwork connecting DNA alkylation and increase in oxidative DNA damage to mutations and structural chromosome aberrations (Fig. 1A) provides a suitable basis for qAOP development. The selected subnetwork draws on two established OECD-endorsed AOPs: AOP 15, which links alkylation of DNA in male pre-meiotic germ cells to heritable mutations (Yauk et al., 2015) and AOP 296 which links oxidative DNA damage to chromosomal aberrations and mutations (Cho, Allemang, et al., 2023). Importantly, multiple *in vitro* assays are available to assess successive KEs along this pathway, including assays quantifying DNA damage, gene mutation, and chromosomal alterations. For some of these assays, an OECD test guideline is available (OECD, 2016a, 2016c, 2016b, 2023). Additionally, alkylating agents comprise a broad class of industrial chemicals, pharmaceuticals, environmental contaminants, highlighting the importance of studying alkylation induced genotoxicity.

**Figure 1.**
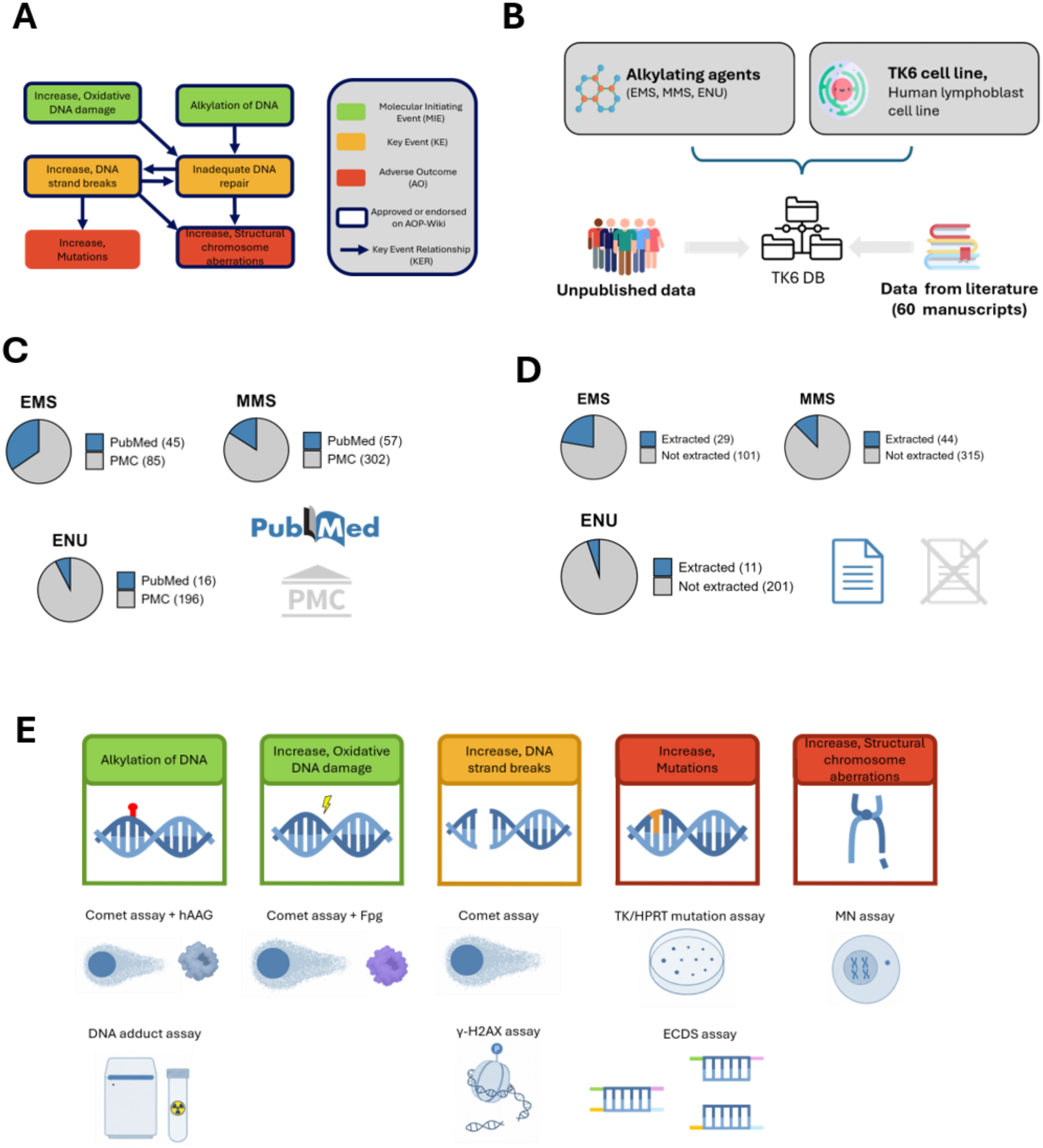
Graphical overview of study design and workflow. [A] The adverse outcome pathway (AOP) subnetwork of interest linking DNA alkylation to increase in mutations and structural chromosome aberrations. [B] A dataset was compiled on quantitative concentration-response data performed on TK6 cells after exposure to either of three alkylating agents: ethyl methanesulfonate (EMS), methyl methanesulfonate (MMS), or N-ethyl-N-nitrosourea (ENU). A total of 63 data sources were utilized for further analysis including 3 unpublished datasets and 60 datasets derived from literature described in independent manuscripts. [C] Number of manuscripts identified through PubMed alongside additional manuscripts identified through PubMed Central (PMC). [D] Number of manuscripts for which data was extracted compared to number of manuscripts deemed irrelevant for data extraction. [E] Overview of the key-events (KEs) in the AOP subnetwork and the representative genotoxic assays for which we had compiled experimental data.

To investigate whether quantitative relationships between the selected KEs can be established that are generalizable across alkylating agents, we first compiled and harmonized a genotoxicity database with data from both publicly available studies and unpublished data on multiple genotoxic endpoints that are relevant for alkylation induced DNA damage. To reduce biological and mechanistic heterogeneity, we restricted our analyses to the human TK6 lymphoblastoid cell line and three prototypical alkylating agents: ethyl methanesulfonate (EMS), methyl methanesulfonate (MMS) and N-ethyl-N-nitrosourea (ENU). Second, we evaluated the extent to which variability between concentration response relationships of different studies can be attributed to experimental design choices. Third, we compared PoDs of genotoxic assays across compounds and experimental designs. Fourth, we quantitatively described KERs within our AOP using response-response modelling.

Overall, our PoD analysis was hampered by large variability both within and between studies, precluding firm conclusions on differences between KEs. Nevertheless, response–response modeling indicated a threshold-like relation in which the upstream KE DNA alkylation tended to take place already at lower concentrations than the downstream KEs DNA strand breaks and chromosome aberrations. Our approach provides a general framework for predicting downstream KE responses from upstream KE responses.

## Methods

### Data collection and systematic literature search

We collected quantitative concentration-response data through a systematic literature search supplemented by unpublished data (Figure 1). For each of the three alkylating agents; EMS (CAS 62-50-0), MMS (CAS 66-27-3), and ENU (CAS 759-73-9), we conducted separate systematic literature searches in PubMed and PubMed Central (PMC). The corresponding search strings and the dates on which these searches were performed are provided in Table S1. We first applied all three search strings to articles indexed in PubMed, searching across all fields, and subsequently to full-text articles available in PMC using the Open Access Subset filter. We focused our analysis on the human TK6 lymphoblastoid cell line, which is widely used in genotoxicity testing. An important limitation of this cell line is its metabolic incompetence. However, this limitation, which can normally be addressed by adding S9 to the culture medium, is not expected to affect the present analysis because the genotoxic potential of EMS, MMS, and ENU does not rely on metabolism (Donovan & Smith, 2010; Klapacz et al., 2016). To determine whether the identified manuscripts were suitable for further analysis, we defined four inclusion criteria: (1) the manuscript must report quantitative data on a genotoxic endpoint that can be linked to a KE within our AOP subnetwork (Fig. 1A); (2) the genotoxic assay of interest must be performed in TK6 cells; (3) the genotoxic assay of interest must involve one of the three selected alkylating agents; and (4) data must be reported for at least two non-zero concentrations of the alkylating agent of interest. We assessed the identified manuscripts in two phases. The first phase involved reviewing abstracts and visually screening figures. We excluded manuscripts that clearly did not meet one or more inclusion criteria at this stage. The final selection was made through full-text evaluation, and we retained only manuscripts meeting all inclusion criteria for further analysis.

### Data extraction

To enable meaningful comparison and analysis of concentration–response data across studies from different sources, we extracted detailed information on experimental design wherever possible. The extracted information was grouped into six main categories, namely: (i) general study descriptors; (ii) factors that may directly influence or bias endpoint measurement; (iii) factors that affect the endpoint indirectly through cytotoxicity; (iv) factors that may affect measurement precision; (v) factors that keep track of units for harmonization of data and (vi) actual measured outcomes (Table S2). To account for heterogeneity in study design, we subdivided the extracted data into groups based on variables with the potential to directly influence or bias endpoint measurement. Whenever available, we obtained raw data or summary statistics that were directly from (supplementary) tables; otherwise, numerical concentration–response values were extracted from figures using a tool for plot digitization, i.e., either PlotDigitizer (*PlotDigitizer*, 2023) or WebPlotDigitizer (Rohatgi, 2017). An overview of all manuscripts from which data extraction was performed can be found in the supplementary materials (Table S3).

### Newly generated data

#### Chemicals, Cells, Culture Conditions

We purchased EMS, MMS and ENU from Sigma Aldrich, St. Louis, MO. We performed all experiments with TK6 cells from ATCC (cat. no. CRL-8015). Details about growth medium and culture conditions can be found elsewhere (S. Avlasevich et al., 2021; Bryce et al., 2016, 2017, 2018).

#### Cell Treatments

Chemical treatments occurred in U-bottom 96-well plates, containing 198 µL TK6 cell suspensions. We added test chemicals, prepared in dimethyl sulfoxide (DMSO), at 2 µL/well resulting in a final DMSO concentration of 1% v/v. We based the top concentration tested on preliminary concentration range-finding experiments, with the goal of achieving approximately 80% reduction to relative nuclei counts (RNC) at 24 hr. In the case of EMS, we tested 19 additional lower concentrations using a square root 2 (i.e., 70.71%) dilution scheme. In the case of ENU and MMS, we included 9 additional lower concentrations, using 70.71% and 50% dilution schemes, respectively. We evaluated each concentration in triplicate wells, and DMSO controls in twelve replicate wells spread throughout the plate. Following addition of test chemicals, an adhesive breathable film and a piece of activated charcoal covered the wells, as described elsewhere (S. L. Avlasevich et al., 2025). Incubation occurred in a humidified atmosphere at 37°C with 5% CO_2_ for 24 hr.

#### MultiFlow DNA Damage Assay: Sample Processing and Flow Cytometric Analysis

We prepared TK6 cells for flow cytometric analysis using reagents and instructions included in the MultiFlow® DNA Damage Kit — p53, γH2AX, Phospho-Histone H3, Cleaved-PARP (Litron Laboratories, Rochester, NY). For components and preparation of the MultiFlow working solution we refer to (Bryce et al., (2016, 2017, 2018). Briefly, at 4 and 24 hr sampling times, we resuspended cells by pipetting, then transferred 25 µL from each well into a new 96-well plate containing 50 µL/well of pre-aliquoted working MultiFlow reagent solution which contained kit-supplied fluorescent antibodies (anti-p53, anti-γH2AX, anti-p-H3, and anti-Cleaved-PARP - note that in our current analysis we only utilized the γH2AX measurements). We performed mixing by pipetting the contents of each well several times. After incubation at room temperature for 30 min, we analyzed samples via flow cytometry using either a Miltenyi Biotec MACSQuant® Analyzer 10 flow cytometer (ENU and MMS experiments), or a FacsCanto II flow cytometer (EMS experiment). We used stock photomultiplier tube detectors and associated optical filter sets to detect fluorescence emissions associated with the fluorochromes Brilliant Violet™ 421, FITC, PE, propidium iodide, and Alexa Fluor® 647. We analyzed flow cytometry data using FlowJo software v10.5.0 (Ashland, OR). We then calculated nuclei to counting bead ratios for each sample, and used these ratios to determine absolute nuclei counts (those with 2n and greater DNA-associated propidium iodide fluorescence). We derived RNC from nuclei counts, and calculated %cytotoxicity as 100 minus %RNC at 24 hr for the MultiFlow assay.

### Quality control and standardization

To improve comparability across studies and reduce technical heterogeneity introduced by differences in reporting formats, we applied extensive quality control and data standardization procedures prior to further analysis. Specifically, we converted reported exposure concentrations to a common unit (µM) using the molecular weights of our test compounds (National Center for Biotechnology Information, 2025b, 2025a, 2025c). Similarly, we harmonized values measured in genotoxicity assays to common reporting scales where possible. For example, we standardized micronucleus frequencies to percentages (%MN), regardless of whether studies originally reported counts per 1000, 2000, or another number of scored cells. When the number of experimental replicates was not explicitly reported, we set replicate counts to values based on the most commonly used replicate numbers within comparable assay types and experimental designs. In ambiguous cases, we used conservative estimates to avoid giving too much weight to these observations.

Furthermore, we standardized reported measures of outcome variability to obtain both standard deviations (SD) and standard errors of the mean (SEM) in a consistent format. In studies reporting interquartile ranges (IQRs) instead of SDs or SEMs, we approximated SDs considering a normal distribution of measurements around the reported mean, in which case the SD is equal to the IQR divided by 1.35. For studies reporting individual replicate-level observations rather than summarized statistics, we merged raw data into summary-level means together with corresponding SDs and SEMs prior to downstream analyses.

To allow for proper comparison of genotoxic responses with different backgrounds, we transformed outcomes to fold-change induction (FC) whenever control measurements were available. Note that this was not possible for DNA adduct data because control responses were not reported in the original study. Moreover, we excluded observations associated with excessive cytotoxicity. Whenever available, we applied a cytotoxicity threshold in accordance with OECD test guideline recommendations. This meant using a cytotoxicity threshold of 50% for the MN and chromosome aberration assay (OECD, 2016a, 2023) and a 90% cytotoxicity threshold for forward gene mutation assays (OECD, 2016b, 2016c). For both the γH2AX assay and the comet assay, we followed the common recommendation for applying a 50% cytotoxicity threshold (Burlinson, 2011; Khoury et al., 2016; Kopp et al., 2019). Since the Error-Corrected Duplex Sequencing (ECDS) assay is relatively new and no formal guidelines or recommendations for cytotoxicity thresholds exist to our knowledge, we used the 90% threshold also applied for the forward gene mutation assay, since both assays test for mutagenicity. When cytotoxicity measurements were unavailable, concentrations exceeding cytotoxicity thresholds observed in comparable studies using similar experimental systems were removed conservatively to minimize potential artefacts.

### Linear mixed effects modeling

To quantify the contribution of experimental design factors to variability in genotoxic concentration–response outcomes, we fitted linear mixed-effects models (LMMs) using the lme4 package in R (Bates et al., 2015) separately for each genotoxic assay. We analyzed FC response values on the natural logarithmic scale, as log-transformation is customary for modeling concentration–response behavior (EFSA Scientific Committee et al., 2022). Because our extracted data reported group means rather than individual observations, direct log-transformation would introduce bias, since the logarithm of a mean is not equivalent to the mean of log-transformed observations (Jensen, 1906; Limpert et al., 2001). Therefore, we bias-corrected log-transformed FC values considering an underlying log-normal distribution of responses. Specifically, we calculated natural log-transformed means according to equation 1 (Limpert et al., 2001):

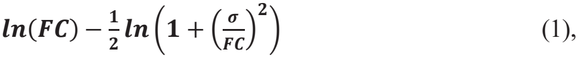

where ***FC*** denotes the reported arithmetic mean of fold-changes and ***σ*** the corresponding SD. For cases where bias correction for the specific observations was not possible due to unavailable variance estimates, we used the median bias-correction term calculated for the remaining observations within that dataset. We modeled concentration–response relationships using a second-degree polynomial for concentration, allowing for non-linear concentration–response behavior that is commonly observed in genotoxicity assays. Models included the experimental design variables identified a priori as potentially influencing endpoint measurement or introducing bias (Table S1) as fixed effects. To account for the hierarchical structure of the compiled dataset, we included random intercepts for concentration–response experiments (hereafter referred to as ‘studies’) nested within published manuscripts, in turn nested within the first author of each manuscript. Missing values in experimental design variables were handled using multiple imputation by chained equations implemented using the mice package in R (Van Buuren & Groothuis-Oudshoorn, 2011).

### Variance decomposition

To estimate the contribution of individual experimental variables to the explained variance of the full mixed effects model, we performed variance decomposition analysis using the partR2 package in R (Stoffel et al., 2021). The procedure has been described in detail elsewhere (Stoffel et al., 2021). Briefly, for each predictor, we refitted a reduced model excluding that predictor while retaining the same random-effects structure. The reduction in marginal R^2^ between the full model and the corresponding reduced model then quantifies the unique contribution of each predictor.

Because predictors could correlate, the sum of uniquely attributed variance across predictors did not necessarily equal the total explained variance of the full model. To account for shared contributions among correlated predictors, we additionally computed inclusive measures of variable importance in the form of inclusive R^2^ values (as implemented in partR2) and Shapley value decomposition (Shapley, 1953) of marginal R^2^. We estimated Shapley values by averaging the marginal contribution of each predictor across all possible combinations of variable inclusion, thereby providing an estimate of each predictor’s overall contribution to explained variance in the presence of multicollinearity. Finally, we quantified variance attributable to hierarchical grouping factors using random-effects variance components extracted from the fitted LMMs. Variance components were expressed as proportions of the total random-effects variance to characterize the relative contribution of between-manuscript, between-author, and between-study variability.

### Data merging

To integrate data across studies, we first combined data pertaining to concentrations that represented effectively the same exposure level. Specifically, concentrations from different studies were considered equivalent when they differed by no more than 1% relative to one another, after which their corresponding response and variability estimates were combined by taking weighted averages of their summary statistics. We selected the study containing the largest number of unique concentrations as the initial reference concentration grid. Concentrations from other studies were then compared against this grid and aligned to an existing concentration when the difference was within the 1% threshold and the match was unambiguous. If no suitable match was found, or if a concentration fell within the 1% threshold of multiple reference concentrations, it was retained as a distinct value and added to the reference grid. This ensured that highly similar exposure levels across datasets were grouped while distinct concentrations were preserved when no clear correspondence existed.

For each merged concentration group, we calculated weighted mean responses and relative survival values using the reported number of replicates as weights (equation 2). We estimated the total variance for each group as the sum of within-study variance and replicate-weighted between-study variance (equations 3-5).

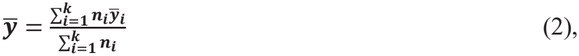

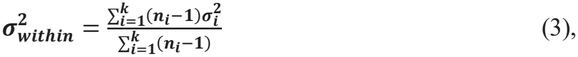

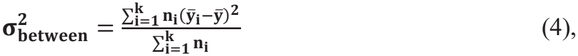

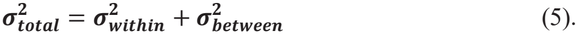

Here, 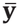 is the overall mean of a given measure across studies and ***k*** is the number of independent studies and ***n***_***i***_ is the number of replicates for study ***i***.

### Benchmark Concentration modeling

We performed benchmark concentration (BMC) modeling (also referred to as benchmark dose / BMD modeling) using a custom R workflow developed to implement the core principles of the PROAST framework (Slob, 2002; Slob & Setzer, 2014) while optimizing the approach for responses expressed as FCs. This custom software has some advantages over existing BMC/BMD tools, specifically the workflow was designed to: 1) accommodate normalized responses with a fixed background response of 1, thereby reducing model complexity, 2) support parallelized computation for the large number of datasets at hand, and 3) provide a flexible basis for extension toward response–response modeling.

Prior to modeling, response means were log-transformed using the same procedure described earlier for the linear mixed-effects modeling. To maintain consistency between the transformed response data and the assumed error structure, we likewise transformed SDs and SEMs to log scale, changing their interpretation from additive to multiplicative (equations 6 and 7) (Limpert et al., 2001):

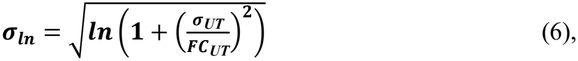

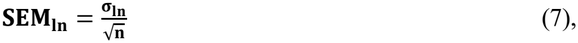

where ***σ***_**In**_ is the estimate for the SD on the natural logarithmic scale, ***σ***_UT_ and ***FC***_***UT***_ the untransformed SD and mean FC respectively and ***n*** is the number of replicates within a study.

We then fitted the log-transformed responses with weights of concentrations equal to the log-transformed SEMs using four families of non-linear growth models: Hill, log-normal, exponential, and Gompertz models (equations 8 – 15 below). For each family, we evaluated both four-parameter (equations 8, 10, 12, 14) and three-parameter (equations 9, 11, 13, 15) variants after parameter optimization to a dataset. In these models, parameter interpretation is broadly consistent across models, with ***b*** controlling the shape and steepness of the concentration-response relationship, ***c*** and ***d*** representing the lower and upper asymptotes respectively and ***e*** acting as a horizontal scaling parameter.

**Hill**

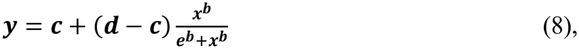

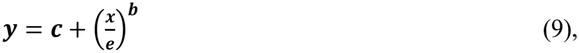

**Log-Normal**

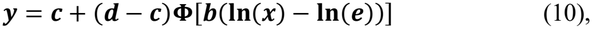

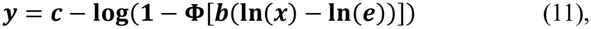

**Exponential**

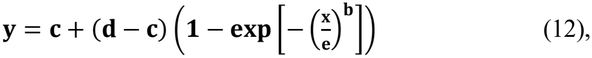

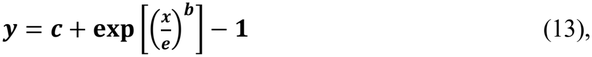

**Gompertz**

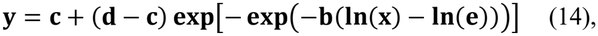

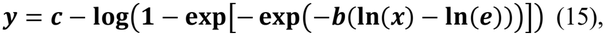

Here, **Φ** is the cumulative distribution function of the standard normal distribution.

For each model family, we selected the model yielding the lowest Akaike Information Criterion (AIC) value and averaged the remaining four models weighing by AIC to obtain an unbiased prediction of genotoxic response at each concentration. This weighting was done based on Akaike weights associated with the AIC differences among the remaining four models (equation 16). Specifically, we here computed the ΔAIC values as the difference between each model’s AIC and the minimum AIC observed among the candidate models. Thus, the best candidate model would have a ΔAIC value of 0 and an associated Akaike weight of 1.

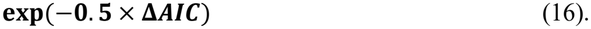

Subsequently, we back-transformed the log-scale weighted predictions to the original response scale. Because back-transformation of log-normal data introduces retransformation bias, we applied a bias correction using concentration-dependent variance estimates. However, observed SD values were only available at experimentally measured concentrations. Therefore, we fitted a simple linear model to the observed untransformed SD values as a function of concentration to obtain continuous proxy SD estimates across the full concentration range. We used these proxy SD estimates both during the initial log transformation procedure and during back-transformation to ensure methodological consistency throughout the workflow. Using the averaged model, we then derived BMCs from the back-transformed weighted prediction curves. The appropriate metric to use as a benchmark response (BMR) within toxicology has been a topic of debate for well over 10 years (Foil et al., 2026; Gollapudi et al., 2013; Slob, 2017; White et al., 2020, 2025; Zeller et al., 2017). Although some have argued that BMR should be evaluated on an endpoint-specific basis (Slob, 2017; Zeller et al., 2017), we chose to use two commonly used types of BMR across all assays instead to allow for a fair comparison of assay sensitivity. Firstly, we chose a BMR of 0.5 corresponding to a 50% increase in the mean control response, which is often used for a variety of *in vitro* genotoxicity tests (Recoules et al., 2024; Wills et al., 2021) and matches the recently recommended BMR for *in vivo* mutagenicity testing (White et al., 2025). This fixed effect size facilitates comparisons across assays, but may not represent an equivalent biological effect across different endpoints. Secondly, we used a BMR corresponding to an increase of one control SD above the mean control response, which is the default value used by the U.S. EPA (U.S. Environmental Protection Agency, 2012). This approach addresses endpoint-specific variability but relies heavily on accurate characterization of background variability.

Finally, we quantified uncertainty in BMC estimates using parametric bootstrapping. For each of the original datasets, we generated 200 bootstrap datasets as follows: For each concentration within a bootstrap iteration, we drew a number of samples equal to the number of replicates in the original dataset at that concentration from a lognormal distribution parameterized using the original mean and SEM values. We then processed each of the resulting bootstrapped datasets through the complete BMC workflow described previously to obtain a range of BMC values from which a 90% CI was acquired.

### Response-response modeling

To quantify KERs, we modeled response-response relationships using paired observations from the merged concentration-response datasets. First, the merged data of two consecutive KEs were matched on nominal concentration. Taking the concentrations of the KE with the most unique concentrations as a reference (or the most upstream KE in case the number of unique concentrations is equal), measurements corresponding to the other KE were considered to correspond to the same nominal concentration when differences in reported concentration relative to one of the concentrations of the reference set were no more than 10%. This resulted in a dataset containing paired responses with both their associated SDs.

Because concentration matching between independent studies often yielded relatively few paired observations, we supplemented the matched dataset with artificial data derived from the BMC bootstrap analyses described above. When fewer than ten matched observations were available, we iteratively selected additional concentrations from the joint concentration range of the two readouts, maximizing their spacing relative to the concentrations already represented in the matched dataset. For example, in the response–response analysis of γH2AX and ECDS for ENU, the joint concentration range spanned 1–150 µM, with approximately overlapping concentrations at 0, 26.5, 53, and 106 µM. The additional concentrations that were selected for collecting artificial data were thus 150 µM, followed by 79.5, 128, 13.25, 39.75, and 92.75 µM. The corresponding bootstrap-derived mean responses and SDs for both KEs were then added to the matched dataset. Specifically, we utilized the distribution of model-averaged bootstrap predictions back transformed to linear scale for each KE to obtain artificial means and SDs on genotoxic responses. This procedure was repeated until at least ten pairs of means and SDs were available for model fitting.

Finally, we fitted the resulting response-response datasets, consisting of a combination of direct observations and bootstrap-based estimates, using the same eight candidate models (from four model families) that were used for concentration-response analysis (Equations 8–15). However, to take into account measurement uncertainty in both readouts, model fitting was performed by orthogonal distance regression (rather than by non-linear regression) as described in detail elsewhere (Boggs et al., 1985; Di Tillio et al., 2026). In brief, the weighted orthogonal distance between the observed data and the fitted model was minimized, with weights equal to the inverse of the variances in both the predictor and response variables. From the 8 resulting model fits, we selected the model yielding the lowest AIC value as the definitive response-response model.

## Results

### Collected genotoxicity data cover the majority of KEs within our AOP subnetwork

To construct a dataset suitable for quantitative comparison of concentration–response relationships across studies, we performed a systematic literature search for genotoxicity data on the alkylating agents EMS, MMS, and ENU (Figure 1). Because cell lines differ in their DNA repair capacity, genomic stability and metabolic competence, which can all influence the outcomes of genotoxic assays (Atha & Reipa, 2023; Khoury et al., 2016), we restricted our literature search to studies conducted in the widely used TK6 cell line. For the γH2AX assay and the MN assay, we supplemented data originating from manuscripts identified through PubMed and PMC with unpublished data (Fig. 1B-D). The resulting dataset contained data on eight distinct genotoxicity assays linked to five of the six KEs described in our AOP-subnetwork, with ‘Inadequate DNA repair’ being the only KE not covered (Fig. 1E, Table S2). Thus, our literature data search delivered an extensive dataset for further quantitative analysis of cellular responses to DNA alkylation.

### Concentration and relative survival are the key factors associated with within-assay variability

Following data extraction, we harmonized all datasets to standardized concentration units, endpoint reporting, measures of variability, and cytotoxicity filtering to maximize comparability across studies before downstream analyses. Despite these efforts to minimize within-assay variability through design of our systematic literature search, the extracted studies varied greatly in terms of their underlying experimental designs (Table S4-S9). This analysis had two purposes: to identify experimental factors that may warrant standardization in future experimental studies and to define relatively homogeneous subsets of data for subsequent meta-analysis.

We utilized two approaches to examine how variability in experimental conditions drives variability in the measured response of the respective genotoxic endpoints. First, we visualized each concentration-response within a given genotoxic assay side-by side and visually separated the data based on experimental conditions (Fig. 2 and S1)). The experimental conditions that best separated groups of concentration-responses differed per assay, which is not surprising given that each genotoxic assay comes with a unique set of choices that can be made for experimental design. One notable factor that impacted concentration-response behavior across genotoxic assays and compounds is time of exposure, where low exposure times typically led to responses only at high concentrations (Fig. 2).

**Figure 2.**
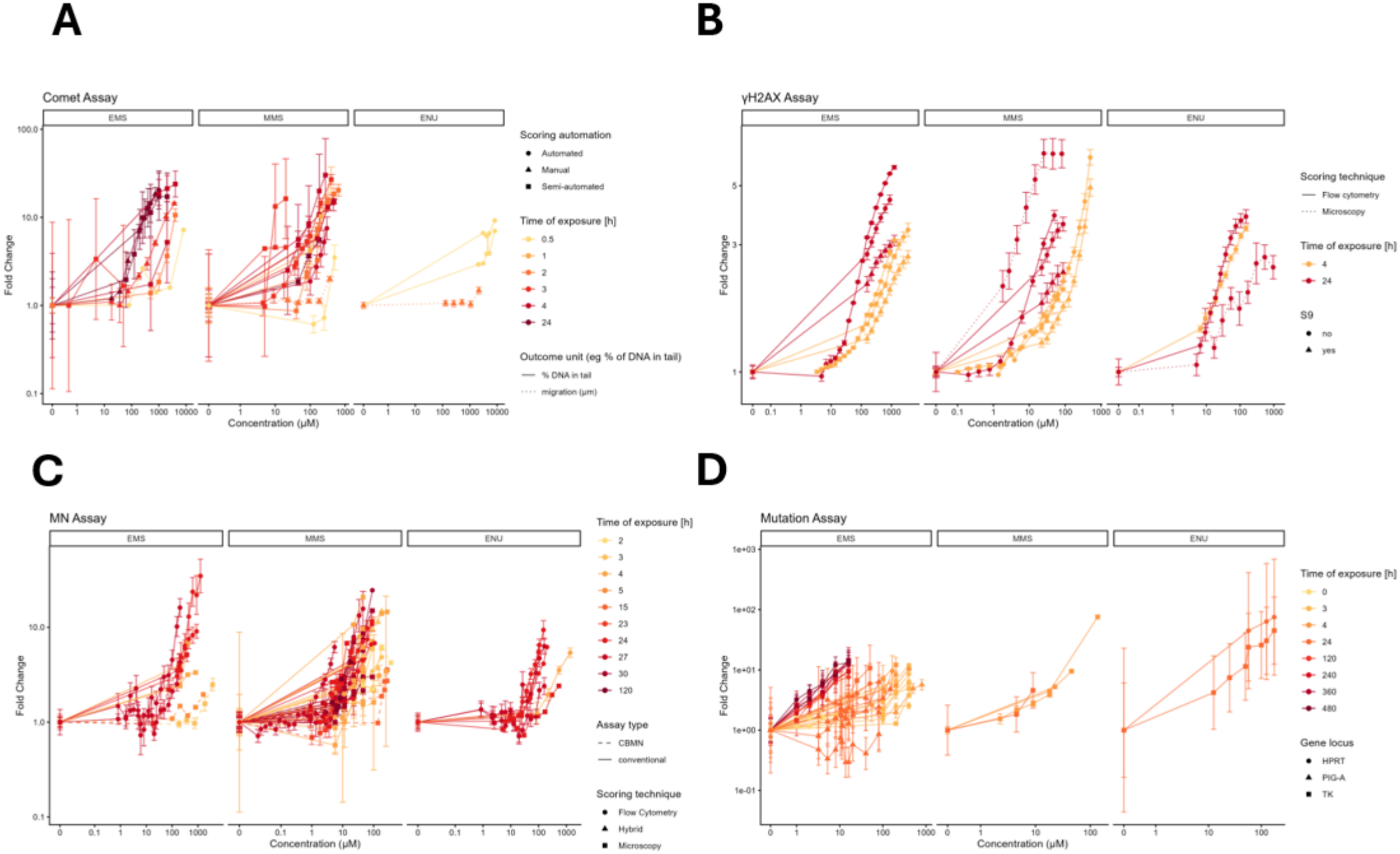
Concentration-response comparisons reveal experimental conditions that drive within-assay variability. Side by side comparisons of concentration-response curves for different experimental set-ups across model compounds. The four genotoxic assays with data available for all three model compounds are depicted: [A] Comet Assay, [B] γH2AX assay, [C] Micronucleus (MN) assay, [D] Gene Mutation assay. Color indicates time of exposure, and symbol shape and line style indicates additional experimental characteristics. Symbols and error bars indicate means and SEMs, respectively.

Second, we performed variance decomposition following linear mixed effects modeling to quantify fixed effects contributions of experimental variables (such as concentration and scoring technique) to the variance explained of the full model. For this approach, we first determined each predictor’s unique contribution to the full model’s R^2^ by iteratively taking out the predictor in question and assessing the reduction in variance explained. From this analysis, concentration emerged as the most important variable explaining a large part of the response variance for comet assays (Fig. 3A, S2A-B); yet, at first sight surprisingly for other assays concentration seemed less important (Fig. 3B-D and S2C-D). For example, for the γ-H2AX assay, relative survival seemed at least as important as nominal concentration (Fig. 3B). However, although this analysis provided insight into the importance of the assessed experimental conditions, the difference in variance explained of the full model and the sum of the uniquely explained variance components was substantial. Thus, this revealed multicollinearity among the predictors which might mask the true importance of the underlying experimental condition. To remedy this, we supplemented our analysis identifying unique variance contributions of individual predictors with importance metrics that aim to better represent the contribution of a predictor including contributions that may (partly) be explained by other predictors, i.e. inclusive R^2^ and Shapley values. From this additional analysis, we found that, as expected, the nominal concentration and the relative survival were for most assays the most influential experimental variables affecting assay output (Fig. 3E).

**Figure 3.**
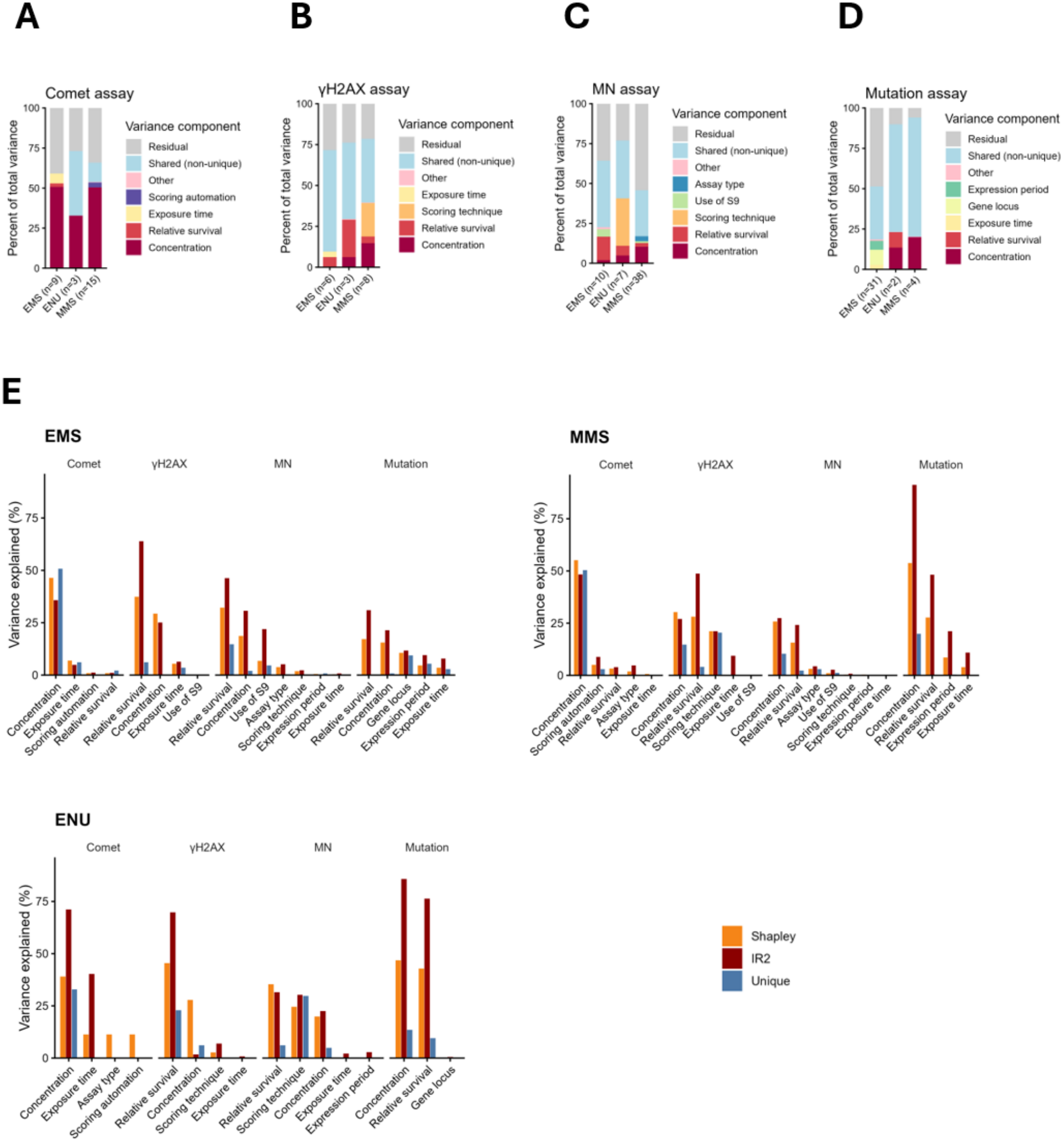
Variance decomposition supports identification of key factors associated with within-assay variability in an unbiased fashion. [A-D] Bars in the color range purple to dark red indicate variance contributions that can uniquely be assigned to a single variable after variance decomposition of a linear mixed effects model in comet assays (A), γH2AX assay (B), Micronucleus (MN) assays (C) and gene mutation assays (D). Variables that individually explain less than 1% of total variance uniquely are grouped together in the ‘other’ category (pink). The shared variance contribution (light blue) is calculated as the difference between the full model’s variance explained and the variance explained by all individual contributors summed up. The residual variance (gray) represents the variance unexplained by the model’s fixed effects. The number of independent studies (n) on which bar plots are based is shown below the bars. [E] Comparison of alternative predictor importance metrics across model compounds. For each experimental variable, bars indicate the percentage of variance attributed based on unique variance explained, inclusive R^2^ (IR^2^), and Shapley value.

In addition to the variance decomposition performed for the fixed effects, we also performed a random-effects variance component analysis to assess the extent to which the remaining variance after accounting for fixed effects could be attributed to the hierarchical structure of study, manuscript and author. This indicated that batch-effects exist at each hierarchical level and that observations are not entirely independent across those groupings (Fig. S2E). In conclusion, the variance decomposition analysis identified nominal concentration and relative survival as the key factors associated with the large variability observed across genotoxicity assays, with a clearly lower yet non-negligible role for other factors.

### BMC modeling does not allow detection of statistical differences in PoDs across KEs

After examining the key factors of variability in our data across compounds and assays, we grouped the collected data into subsets that are similar in experimental design for those key factors. We assigned each subset a standardized code representing their design in terms of the identified key experimental factors. The code consists of three or four letters, depending on the assay type (Table S4-S9, Fig. 4A). As an example: the code SEMB is used to indicate the MN data pertaining to short exposure (indicated by ‘S’), early harvest (indicated by ‘E’), microscopic scoring (indicated by ‘M’), and using the CBMN assay format (indicated by ‘B’). To increase concentration coverage and improve precision, we merged data from independent studies but sharing the same subgroup code for further analysis while accounting for both within-study and between-study variability (Fig. S3A).

**Figure 4.**
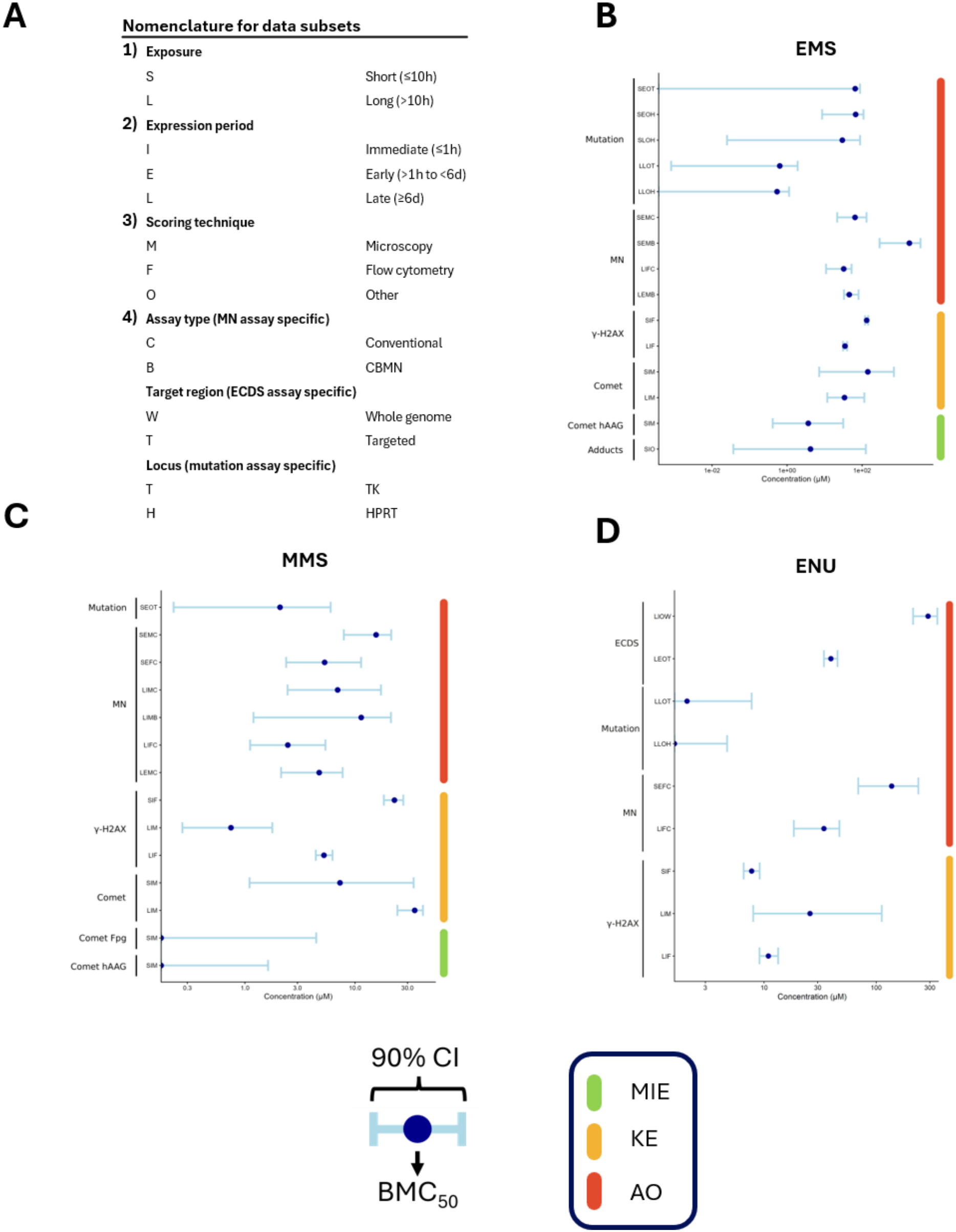
Benchmark concentration modeling AOs does not allow detection of statistical differences in PoDs across KEs. [A] Nomenclature for subsets based on experimental design. Eg: SIO stands for Short exposure, immediate harvest and a scoring technique other than microscopy or flow cytometry. [B-D] Benchmark concentration associated with 50% increase relative to control (BMC_50_; dots) values alongside 90% CIs (error bars) across genotoxic assays and experimental conditions. Bootstrap estimates were calculated from 200 bootstrap datasets. The first three panels represent our three model compounds ethyl methanesulfonate (A), methyl methanesulfonate (B), or N-ethyl-N-nitrosourea (C).

We performed BMC modeling on each resulting merged dataset to compare PoDs across assays and compounds (Fig. S3B-D). To estimate uncertainty around the BMC estimates, we generated 90% confidence intervals by resampling the data in log space from a lognormal distribution.

Overall, the BMC confidence intervals were often broad, reflecting strong variability in the BMC estimates for different bootstrap samples of the underlying data (Fig. 4, Fig. S4, Table S10). The width of the confidence intervals varied across assays and subsets, with narrower intervals generally observed for the more recently developed γH2AX and ECDS assays, and for subsets consisting of fewer datasets from different sources (Table S10). These observations suggest that insufficient concentration coverage of older studies and strong heterogeneity among studies with similar experimental design contribute to the uncertainty in the estimated PoDs.

Despite this uncertainty, the BMC estimates showed a general tendency towards lower PoDs for assays linked to the MIEs ‘Increase, Oxidative DNA damage’ and ‘Alkylation of DNA’ than for assays associated with the downstream KE ‘Increase, DNA strand breaks’ and the AO ‘Increase, Structural chromosome aberrations’ (Fig. 4, Fig. S4, Table S10). This tendency is consistent with the expected ordering of events in an AOP, in which MIEs precede downstream KEs and AOs. However, the broad and often overlapping confidence intervals preclude strong conclusions regarding differences in sensitivity between these events. The forward gene mutation assay formed an exception to this tendency, i.e. the BMC estimates for this assay were generally lower (although often not statistically significant) compared to assays representing more upstream KEs. However, this apparent discrepancy may reflect differences in study design and assay characteristics. The available forward mutation datasets largely originate from studies conducted in the 1980s and 1990s that were not designed with BMC modeling in mind. Moreover, the prolonged expression periods required to allow mutations to be formed, distinguish forward gene mutation assays from most other genotoxicity assays and complicate comparisons of their estimated PoDs.

In conclusion, exposure to DNA alkylating agents in general led to triggering MIEs at somewhat lower concentrations than for the intermediate KEs and AOs, although variability of the data precluded strong conclusions.

### Response-response modeling allows for prediction of late-stage downstream KEs from upstream KEs

To establish qKERs, we derived response-response relationships for adjacent and non-adjacent KEs for our EMS and MMS datasets. These relationships were based on paired observations obtained by matching concentration-response data from the corresponding KEs at the same nominal concentration (Fig. 5 and S5, red points). Because relatively few matched observations were available for most KERs, the datasets were supplemented with artificial data generated from the bootstrap model predictions that were used in our BMC analysis (Fig. 5 and S5, black points).

**Figure 5.**
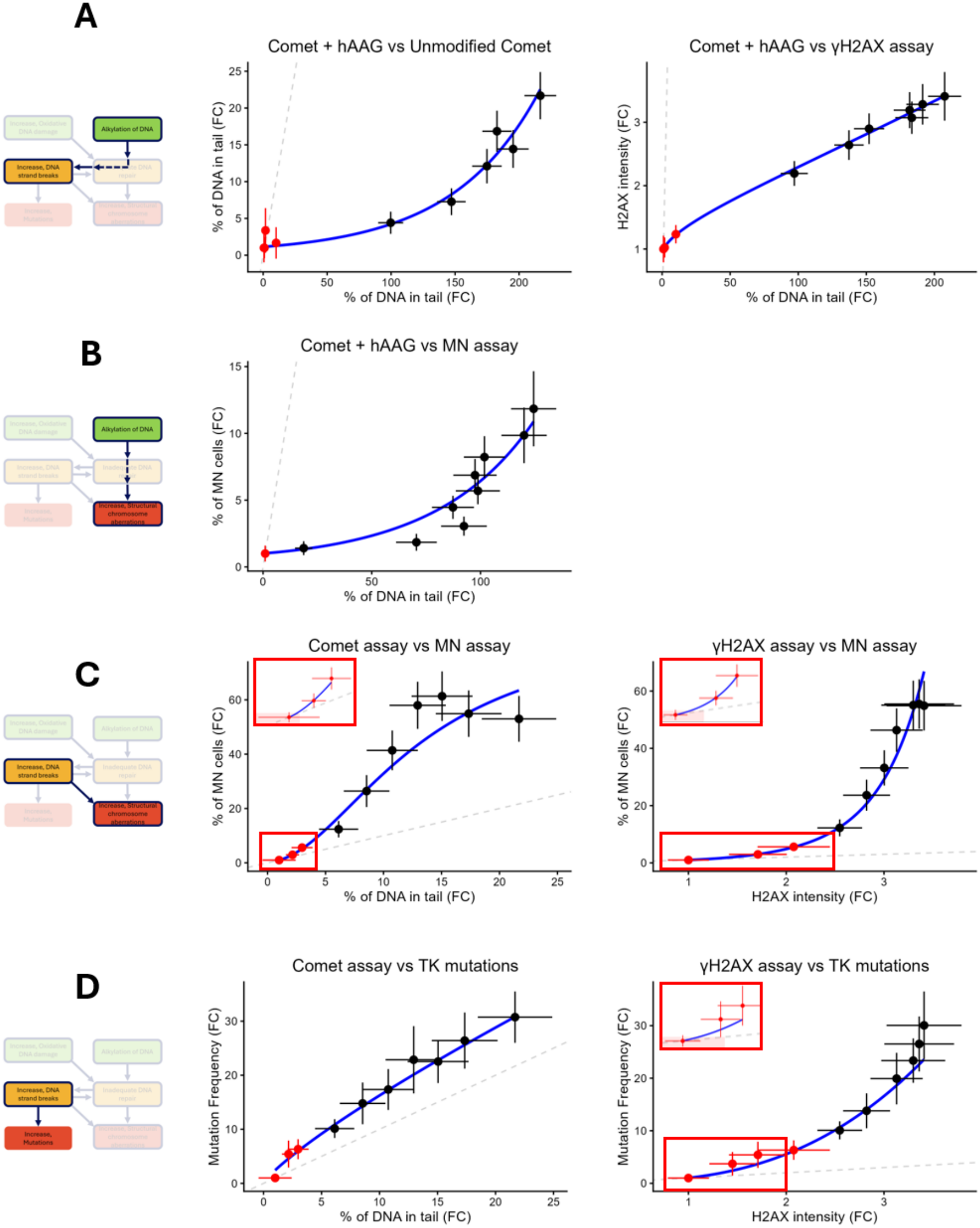
Response-response modeling for EMS allows for prediction of late stage KEs from upstream KEs. [A-D] Fitted response-response models for the following adjacent and non-adjacent KEs: Alkylation of DNA leads to Increase in DNA strand breaks (A), Alkylation of DNA leads to increase in structural chromosome aberrations (B), DNA strand breaks lead to increase in structural chromosome aberrations (C) and DNA strand breaks lead to increase in mutations (D). Responses on all genotoxic endpoints are reported as fold changes (FCs) relative to control. Red dots indicate experimentally matched pairs of mean responses whereas black dots indicate artificial data in the form of means obtained from bootstrap prediction curves through benchmark concentration (BMC) modelling. Error bars indicate the square root of SDs on either observed data (red) or artificial data (black). The leftmost panels indicate the studied key-event relationships (KERs), where solid lines are used for adjacent KERs and dashed lines for non-adjacent KERs. In the middle and right panels, the grey dashed line indicates the identity line (y=x). Insets within red boxes show zoom-ins at low responses, with red shaded rectangles denoting the 1.5-fold responses (corresponding with the BMRs).

Subsequently, we fitted response-response models to these combined datasets (Fig. 5 and S5, blue lines). For each of the KERs examined, we established a well-fitting response-response model (Table S11) on either of eight non-linear growth models from four families (equations 8-15). Interestingly, the relationship between DNA adducts measured using the hAAG-modified comet assay and DNA strand breaks as measured through the γH2AX assay very closely approximated linearity for EMS and MMS (Fig. 5A, Fig. S5A). However, most response-response relationships are highly nonlinear and suggest threshold like relationships between several KEs. For the KERs relating to the MIE of DNA alkylation, the initial slope of the response-response models is much smaller than 1 (Fig. 5A-B, Fig. S5A-B), which indicates that the more downstream KE or AO is triggered at lower concentrations than the MIE. For subsequent KERs, the initial slope of the response-response curves is much closer to 1 (Fig. 5C-D and S5C-D), indicating that the difference between the responses at low concentrations is less apparent. Note that the response-response relations for EMS (Fig. 5) and MMS (Fig. S5) are similar in shape but can quantitatively differ by up to approximately 2-fold. Response–response relationships for ENU (Fig. S6) generally followed the same trend as those for EMS and MMS. However, a direct comparison cannot be made because short-exposure data, as used in the EMS and MMS analyses, were not available for all KEs in the ENU dataset. In conclusion, the fitted response-response models allow to predict responses associated with later KEs from responses associated to earlier ones. This response-response analysis suggests that DNA adduct formation as measured through the hAAG modified comet assay is triggered at lower concentrations compared to assays associated with more downstream KEs.

## Discussion

In this study, we provided a proof of concept for the quantitative development of an AOP for genotoxicity using DNA alkylation as a model case. We leveraged the extensive body of existing *in vitro* genotoxicity data to assess to what extent these data can be integrated into a quantitative framework and to identify remaining data gaps. We collected a harmonized dataset of concentration–response data from TK6 cells exposed to either of three alkylating agents (EMS, MMS and ENU), encompassing eight genotoxicity assays representing five biological events in the AOP (MIEs, KEs or AOs), together with extensive information on experimental design per data subset. These data enabled us to compare the *in vitro* PoDs of genotoxic endpoints across assays and experimental designs, and to establish qKERs linking upstream to downstream KEs. Collectively, our results provide insight into both the opportunities and challenges associated with quantitatively integrating heterogeneous concentration–response data.

The development of qAOPs has increasingly been proposed as a means of translating the mechanistic structure of AOPs into quantitative predictions for hazard and risk assessment (Conolly et al., 2017; OECD, 2018; Perkins et al., 2019). Mechanistic systems-toxicology approaches may be most promising for general risk assessment due to their ability to describe the temporal dynamics of KEs and their interactions and therefore offer the greatest potential to capture the dynamic nature of an AOP (Di Tillio & Beltman, 2024). However, such models require sufficiently detailed time-course data on multiple KEs, which for genotoxicity is virtually nonexistent. Moreover, genotoxicity data have traditionally been generated primarily for qualitative hazard identification rather than quantitative analysis (Menz et al., 2023). Indeed, existing qAOP approaches have generally been limited by the availability and quality of quantitative data needed to parameterize and validate the models (Jeong et al., 2025; Spinu et al., 2020). Within this context, response–response modeling provided a pragmatic approach for our proof-of-concept study. It allowed us to establish qKERs using the wide range of concentration–response data available in the literature.

The BMC estimates obtained from our merged datasets with comparable experimental designs were broadly consistent with estimates reported for some of the underlying studies. Out of the 60 manuscripts from which data were extracted, six manuscripts reported BMC values (Table S12) (Bemis et al., 2016; Cho, Swartz, et al., 2023; Rees et al., 2017; Sanders et al., 2022; Verma et al., 2017, 2018). These six studies used BMRs ranging from a 5% to a 400% increase over control, and all used some version of PROAST for BMC modeling. Overall, the reported BMC_50_ estimates were in good agreement with those obtained in our analysis for the corresponding subgroups. For example, for the ECDS subgroup LEOT (referring to long exposure (L), early harvest (E), response measurement other than flow-cytometry / microscopy based (O), for a targeted genomic region (T)), our analysis yielded a 90% BMC_50_ CI for ENU of 34.4–45.2 µM, compared with average lower and upper limits of the CI being 33.3 and 47.8 µM, described respectively based on BMC analysis of the underlying data in PROAST (Table S12) (Cho, Swartz, et al., 2023). These experiments represented two laboratories and three harvest times, with BMC analysis performed separately for each experiment in the source study. Similarly, for the MN assay subgroup LEMB, our calculated 90% BMC_50_ CI for EMS (32.9–80.4 µM) was comparable to the BMC_50_ 90% CI 36.3– 99.1 µM reported in literature for the underlying data (Sanders et al., 2022). In many other cases, however, the BMCs calculated in our analysis represent concentration–response behavior across multiple laboratories and therefore incorporated between-study heterogeneity that was not captured by BMC estimates derived from individual studies. Consequently, our confidence intervals were often wider than those reported for individual experiments. For example, literature reported a 90% BMC_50_ CI of 919.1–2004.4 µM for 3h exposure MN data after exposure to EMS (Sanders et al., 2022), whereas analysis of the corresponding combined subgroup in our study (SEMB) yielded a substantially wider CI of 292.8– 3604.1 µM.

An important finding in this study is that many existing concentration–response data do not provide sufficient information for reliable estimation of PoDs. Although the BMC analysis suggested that the MIEs of DNA alkylation and increase in oxidative DNA damage generally occurred at lower concentrations than downstream KEs, the confidence intervals around many BMC estimates were wide and often overlapping, preventing a clear distinction in sensitivity between genotoxic assays. This uncertainty stems from two distinct issues: First insufficient characterization of the concentration–response relationship and secondly, heterogeneity between studies from different sources that we merged for BMC analysis yet reflected the same endpoint and general experimental set-up. With respect to the first point, reliable estimation of a BMC requires sufficient information to characterize a concentration-response relationship. Data spanning low, intermediate, and high response levels are important, with observations at concentrations relating to low response levels being particularly pertinent because they best capture the concentration-response behavior around the BMR/BMC (EFSA Scientific Committee et al., 2022). When the lowest tested non-zero concentration already produces a response substantially exceeding the BMR, or when the highest tested concentration does not reach the BMR, a wide range of curves may explain the available data, resulting in imprecise BMC estimates. Our findings thus underscore the importance of sufficient concentration coverage for concentration-response studies intended for quantitative analysis and reinforce existing recommendations (EFSA Scientific Committee et al., 2022). Many existing genotoxicity studies were primarily designed for hazard identification but provide insufficient data for quantitative PoD estimation even when combined with data from similar studies. Future studies intended to estimate PoDs and/or qAOP development should therefore use sufficiently broad and well-spaced concentration ranges that capture low, intermediate, and high response levels, preferably using at least three biological replicates per concentration. With respect to the second point, the uncertainty in PoD estimation also reflects substantial heterogeneity in concentration–response behavior between studies. Even after accounting for differences in assay and test compound, considerable variability in genotoxic responses remained between studies. Concentration and relative survival accounted for a substantial proportion of this variability in the variance decomposition, although these contributions cannot be directly interpreted as causal effects. Nevertheless, a causal interpretation of the association with nominal concentration is biologically plausible, given that concentration determines the amount of test compound to which cells are exposed. In contrast, relative survival is itself a measured response and is likely to reflect downstream effects, although in some cases it may capture effective exposure better than nominal concentration. For example, effective exposure can be reduced by binding of compound to well-plate plastic, cellular metabolism, abiotic degradation (Henneberger et al., 2025) or lack of membrane transporters (Ciarimboli, 2012). Differences in how relative survival was measured may however weaken this association, potentially explaining why nominal concentration accounted for the largest proportion of variance in some analyses whereas relative survival did so in others. We expected the effect of relative survival to be stronger for the gene mutation assay, where we applied a more lenient cytotoxicity threshold compared to the other assays (90% vs 50%), however, our analysis indicated no clear difference in the importance of relative survival between the gene mutation assay and other assays like the MN assay and γH2AX assay.

Importantly, the contributions of individual experimental design characteristics should also be interpreted in the context of the available data. A predictor can only account for variance when sufficient variation in that predictor exists among the studies included in the analysis. Thus, a lack of estimated contribution does not necessarily imply that an experimental condition does not have the potential to influence a response. For example, we could not use variance decomposition to identify whether prolonged expression period potentially affected comet assay responses because no comet assay datasets in the compiled data included such an expression period. In addition, correlations between experimental design variables make it difficult to uniquely attribute shared variance to individual predictors. Our variance decomposition should therefore be viewed as an attempt to identify factors associated with differences in response among the available studies, rather than as establishing the causal importance of all possible individual experimental conditions.

Importantly, only a small part of the between-study heterogeneity could be associated with experimental design characteristics beyond concentration and relative survival. Variance component analysis did identify batch effects at multiple levels of hierarchy within the data: between measurements within concentration–response experiments, between experiments originating from the same manuscript, and between experiments conducted by the same researcher (assuming that the first author represents the researcher conducting the experiments). These random effects together accounted for approximately one third of the variance attributable to the random-effects structure. All in all, our variance decomposition and variance component analyses suggest that differences between publicly available studies cannot be fully reconciled by accounting for the experimental design characteristics reported in these studies. For future qAOP development, generating data for multiple KEs within the same laboratory using similar experimental procedures would therefore be advantageous, as this will at least reduce the heterogeneity arising from laboratory- and study-specific factors, facilitating reliable quantitative integration across KEs.

Despite substantial heterogeneity and data gaps, we demonstrated that existing concentration–response data can be used to establish quantitative relationships between KEs with response-response modeling. This analysis supported the conclusion that DNA alkylation, as the MIE, may occur at lower concentrations than subsequent KEs, while the available data did not allow the order of the downstream KEs to be distinguished with confidence. This observation could be due to the similar level of biological organization (i.e., cellular level) at which these measured key events all take place.

Importantly, the established qKERs also provide a quantitative framework for linking responses across KEs rather than considering each genotoxicity endpoint independently. Such relationships may enable prediction of downstream KEs from upstream responses when measurements are unavailable, thereby providing a potential means of filling data gaps in quantitative risk assessment. Quantitative relationships between KERs are similar across EMS and MMS up to a fold difference of approximately 2. For instance, a 10-fold increase in comet assay response is associated with an approximately 30-fold increase in MN assay response upon EMS exposure, but with an approximately 15-fold increase upon MMS exposure (compare Fig. 5C with S5C, left panels). This finding is consistent with earlier work showing that downstream responses can substantially differ after exposure to compounds with similar modes of action despite similar upstream responses (Burgers et al., 2026). This suggests that additional mechanistic understanding is required of the processes leading to genotoxic events before these response-response relations can be quantitatively extrapolated to other compounds.

The general framework developed here is not limited to DNA alkylation or genotoxicity. When sufficiently informative concentration–response data are available for the relevant KEs, response–response modeling can be used to quantify relationships between biological events and may allow unmeasured downstream responses to be estimated. This approach offers a way to integrate heterogeneous experimental data into qAOPs and could be extended to other AOPs and toxicological pathways. Overall, our findings indicate that existing genotoxicity data can support qAOP development, but that the reliability of quantitative integration is strongly dependent on the design and consistency of the underlying concentration–response studies. Designing future studies with quantitative integration in mind, while making use of response– response relationships in existing data, is therefore needed to enable robust and predictive qAOPs.

## Statements and Declarations

### Code and data availability

All code and original data required for the analyses presented in this manuscript will be made publicly available on GitHub and Zenodo upon publication.

### Ethics approval

No approval of research ethics committees was required to accomplish the goals of this study because experimental work did not involve animal experiments.

## Author contributions

L.W.: Conceptualization, software writing, literature search, data extraction, data analysis, validation, visualization, writing - original draft, writing - review & editing; E.D.: literature search, data extraction, writing - review & editing; S.B.: Performance of wet-lab experiments, data curation, writing - review & editing; M.S.: Data extraction, writing - review & editing; A.T.: Data extraction, writing - review & editing; F.G.: Data extraction, writing - review & editing; S.M.: Data extraction, writing - methods for new experimental data, writing - review & editing; T.P.: Literature search, writing methods section on literature search, writing - review & editing; M.A.: Conceptualization, supervision, funding acquisition, writing - review & editing; B.M.: Conceptualization, data extraction, supervision, funding acquisition, writing - review & editing; J.B.: Conceptualization, supervision, funding acquisition, writing - review & editing.

