## Supplementary Figures for "Advancing AOPs for genotoxicity through quantifying KERs linking DNA alkylation to increase in mutations and chromosome aberrations"

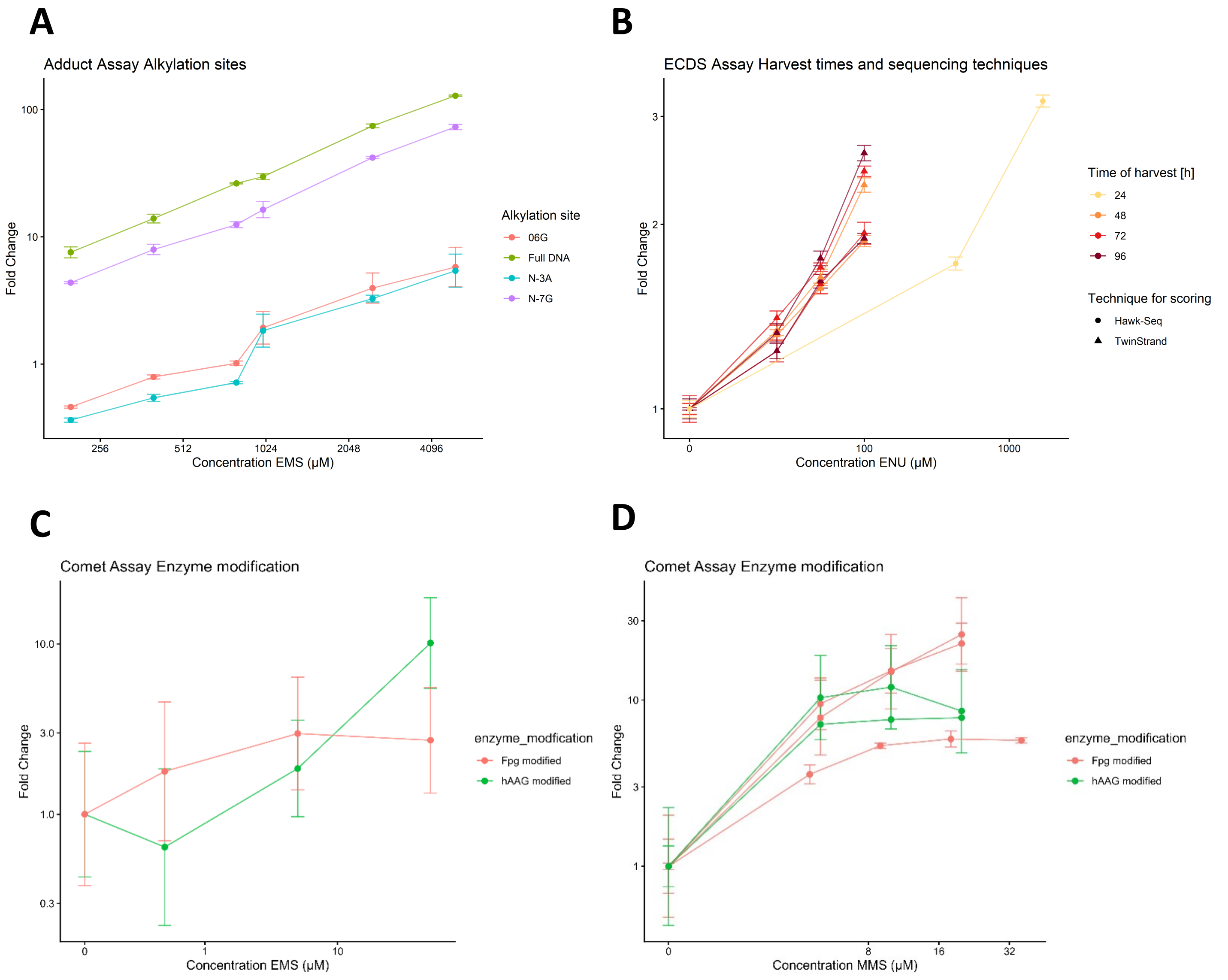


**Figure S1: Concentration-response comparisons of extracted data.** [A-D] Side-by-side comparisons of concentration–response curves for different experimental set-ups. Shown are the genotoxic assays that did not contain data for all three model compounds to complement main text Figure 2. Depicted assays include: DNA adduct assay (A), ECDS assay (B), Fpg and hAAG modified comet assays for EMS (C) and MMS (D).

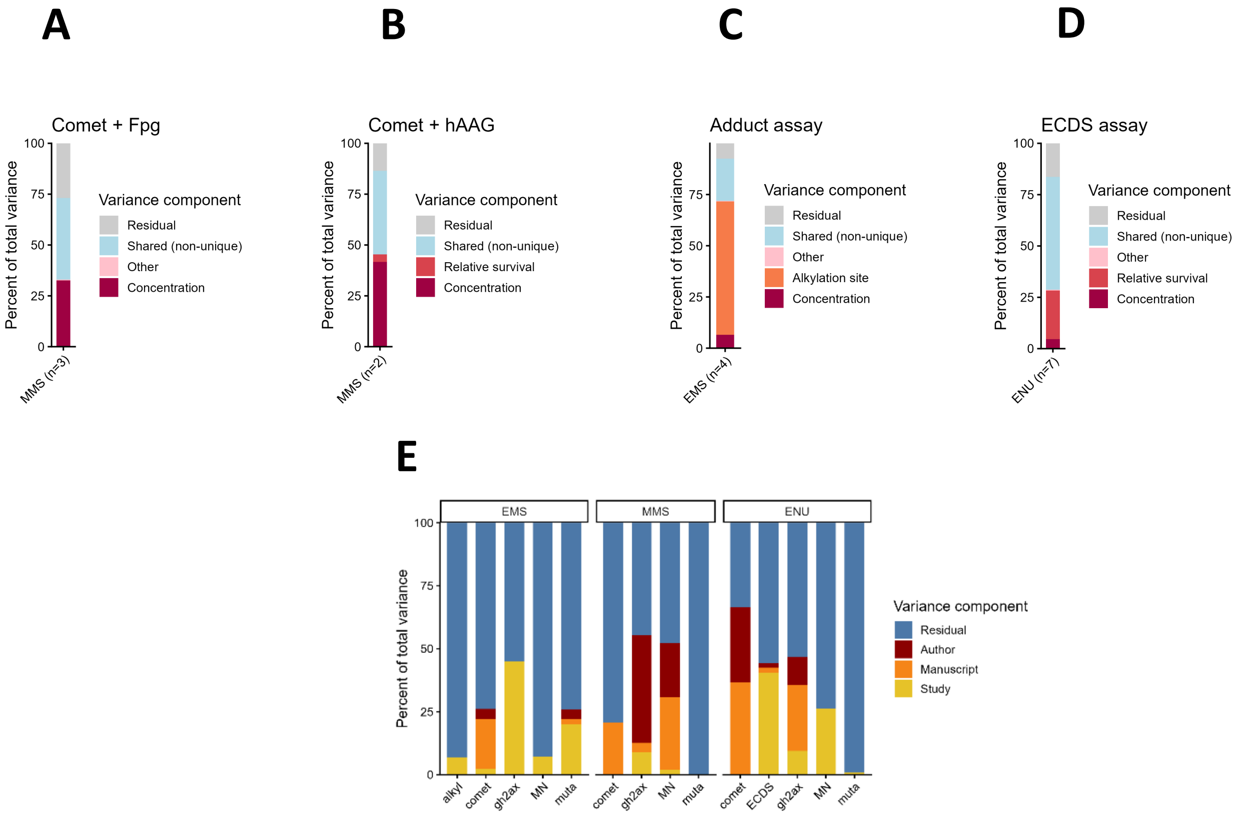


**Figure S2: Variance decomposition and variance component analysis.** [A-D] Bars in the color range purple to dark red indicate variance contributions that can uniquely be assigned to a single variable after variance decomposition of a linear mixed effects model in Fpg-modified comet assays (A), hAAG-modified comet assays (B), DNA adduct assay (C) and ECDS assay (D). Variables that individually explain less than 1% of total variance uniquely are grouped together int the ‘other’ category (pink). The shared variance contribution (light blue) is calculated as the difference between the full model’s variance explained and the variance explained by all individual contributors summed up. The residual variance (gray) represents the variance unexplained by the model’s fixed effects. [E] Variance component analysis of random effects from the fitted linear mixed-effects models. Bars indicate the variance assigned by the model to each of the random effects.

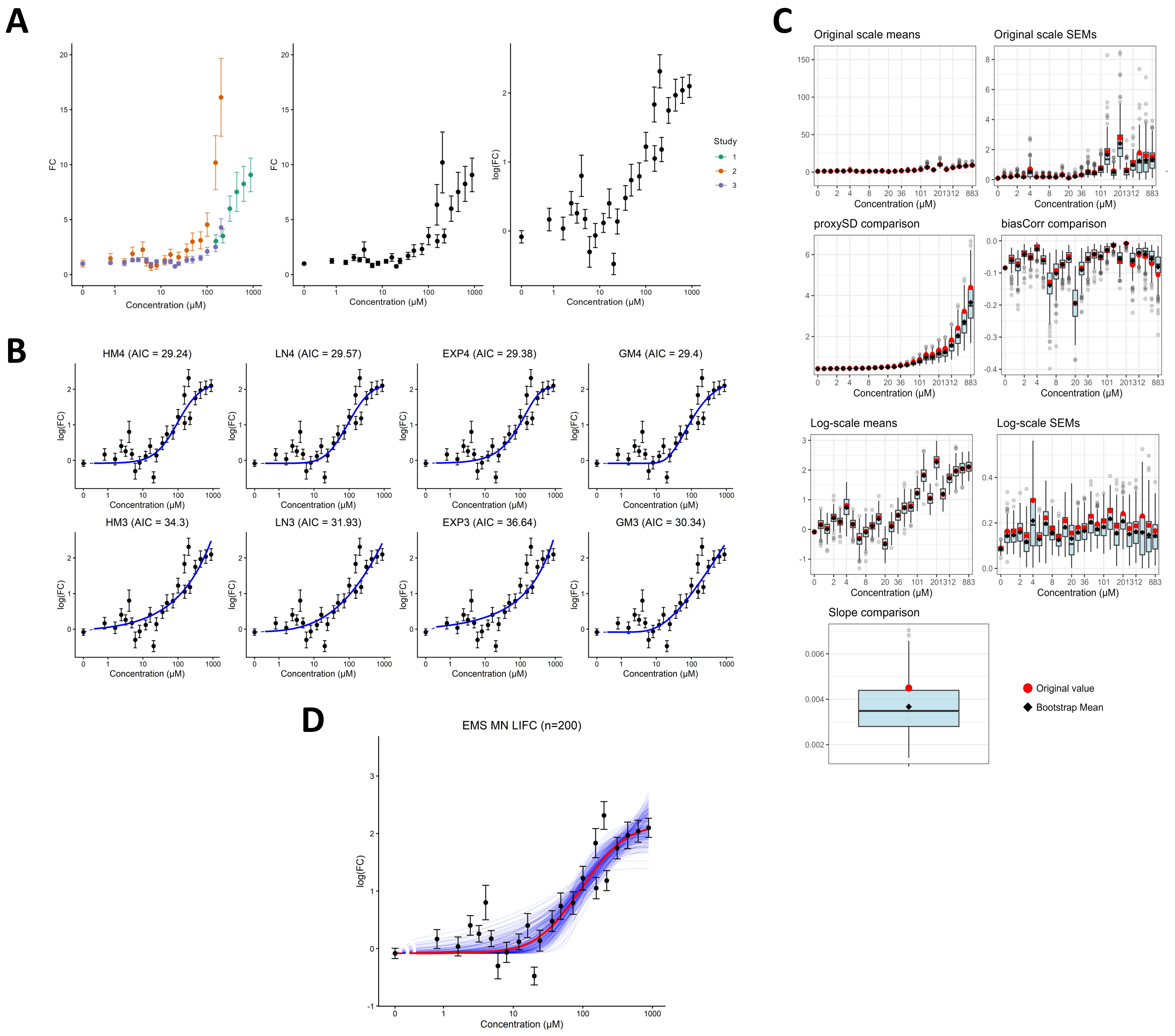
A
B
C
D
FC
log(FC)
Concentration (μM)
Study
1
2
3
HM4 (AIC = 29.24)
LN4 (AIC = 29.57)
EXP4 (AIC = 29.38)
GM4 (AIC = 29.4)
HM3 (AIC = 34.3)
LN3 (AIC = 31.93)
EXP3 (AIC = 36.64)
GM3 (AIC = 30.34)
Original scale means
Original scale SEMs
proxySD comparison
biasCorr comparison
Log-scale means
Log-scale SEMs
Slope comparison
Original value
Bootstrap Mean
EMS MN LIFC (n=200)

**Figure S3: Workflow for BMC and CI estimation applied to studies sharing common experimental design.** [A] Data preparation and merging. Concentration-response data from individual studies using comparable experimental design (left panel) are merged at identical concentrations to generate a single merged dataset (middle panel), and merged responses are log-transformed before subsequent concentration-response modeling (right panel). [B] Model fitting and model averaging. The log-transformed merged dataset is fitted using four concentration-response model families, each represented by a three- and a four-parameter variant (effectively two and three parameters because c is fixed to 1), resulting in eight candidate models. Model fits are subsequently used for model averaging and BMC estimation. [C] Parametric bootstrap procedure. Distributions of key model inputs and parameters obtained from 200 bootstrap datasets are shown, including response means, standard errors of the mean (SEMs), proxy standard deviations (proxySDs), bias-correction terms applied during log- (back)transformation, and the intercept and slope parameters of the proxySD model. [D] Bootstrap-derived concentration-response fits. Concentration-response curves obtained from the 200 bootstrap datasets (blue) are shown together with the fit to the original dataset (red) and the original log-transformed observations. The distribution of bootstrap fits is used to quantify uncertainty in the BMC estimate and derive the associated 90% confidence interval.

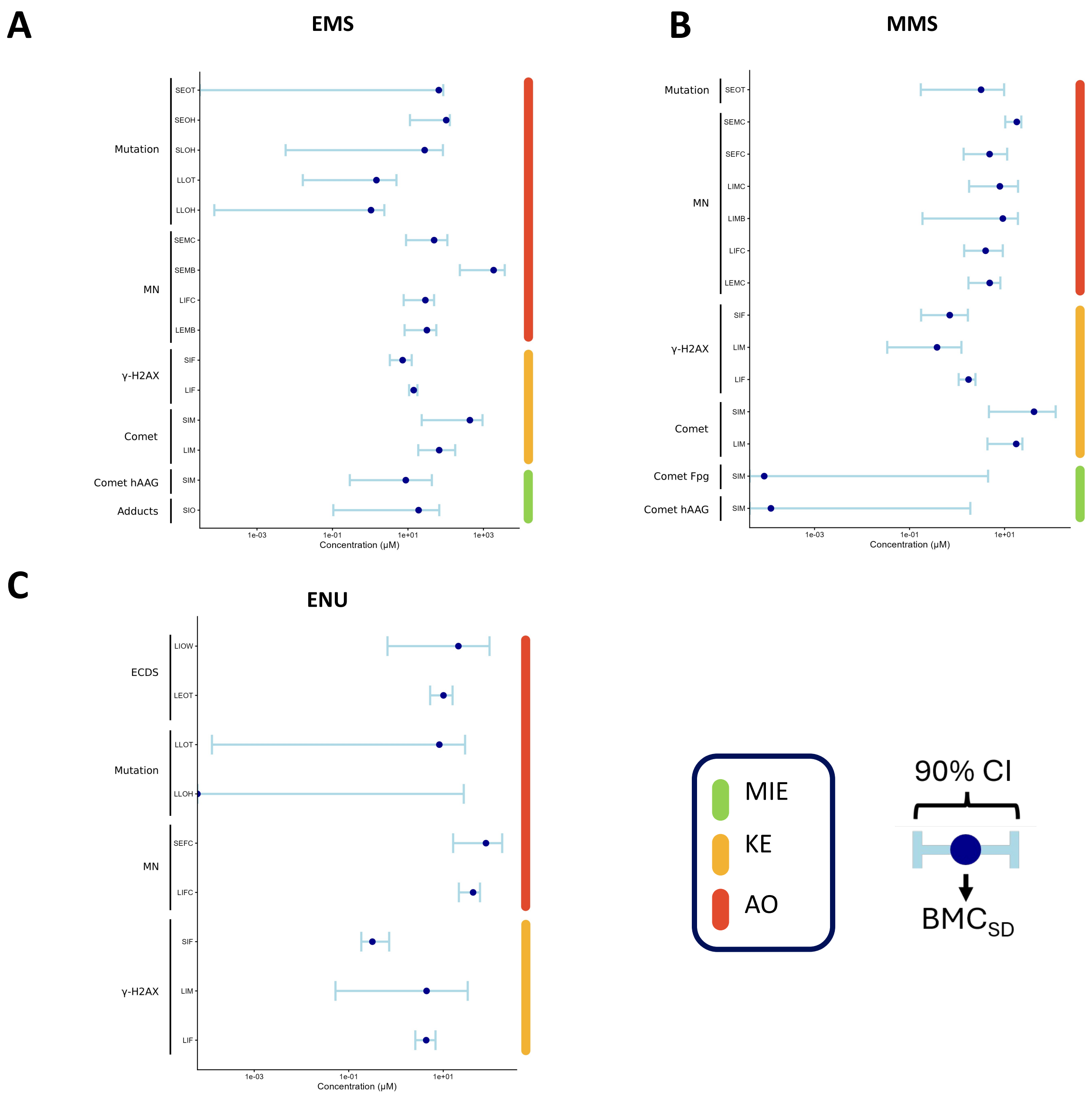

A
EMS
Mutation
SEOT
SEOH
SLOH
LLOT
LLOH
MN
SEMC
SEMB
LIFC
LEMB
γ-H2AX
SIF
LIF
Comet
SIM
LIM
Comet hAAG
SIM
Adducts
SIO
1e-03
1e-01
1e+01
1e+03
Concentration (μM)
B
MMS
Mutation
SEOT
MN
SEMC
SEFC
LIMC
LIMB
LIFC
LEMC
γ-H2AX
SIF
LIM
LIF
Comet
SIM
LIM
Comet Fpg
SIM
Comet hAAG
SIM
1e-03
1e-01
1e+01
Concentration (μM)
C
ENU
ECDS
LIOW
LEOT
Mutation
LLOT
LLOH
MN
SEFC
LIFC
γ-H2AX
SIF
LIM
LIF
1e-03
1e-01
1e+01
Concentration (μM)
MIE
KE
AO
90% CI
BMCSD

**Figure S4: Benchmark concentration modeling Using $BMC_{SD}$.** BMCSD values (dots) alongside 90% CIs (error bars) across genotoxic assays and experimental conditions for EMS (A), MMS (B) and ENU (C). Bootstrap estimates were calculated from 200 randomly sampled bootstrap data frames.

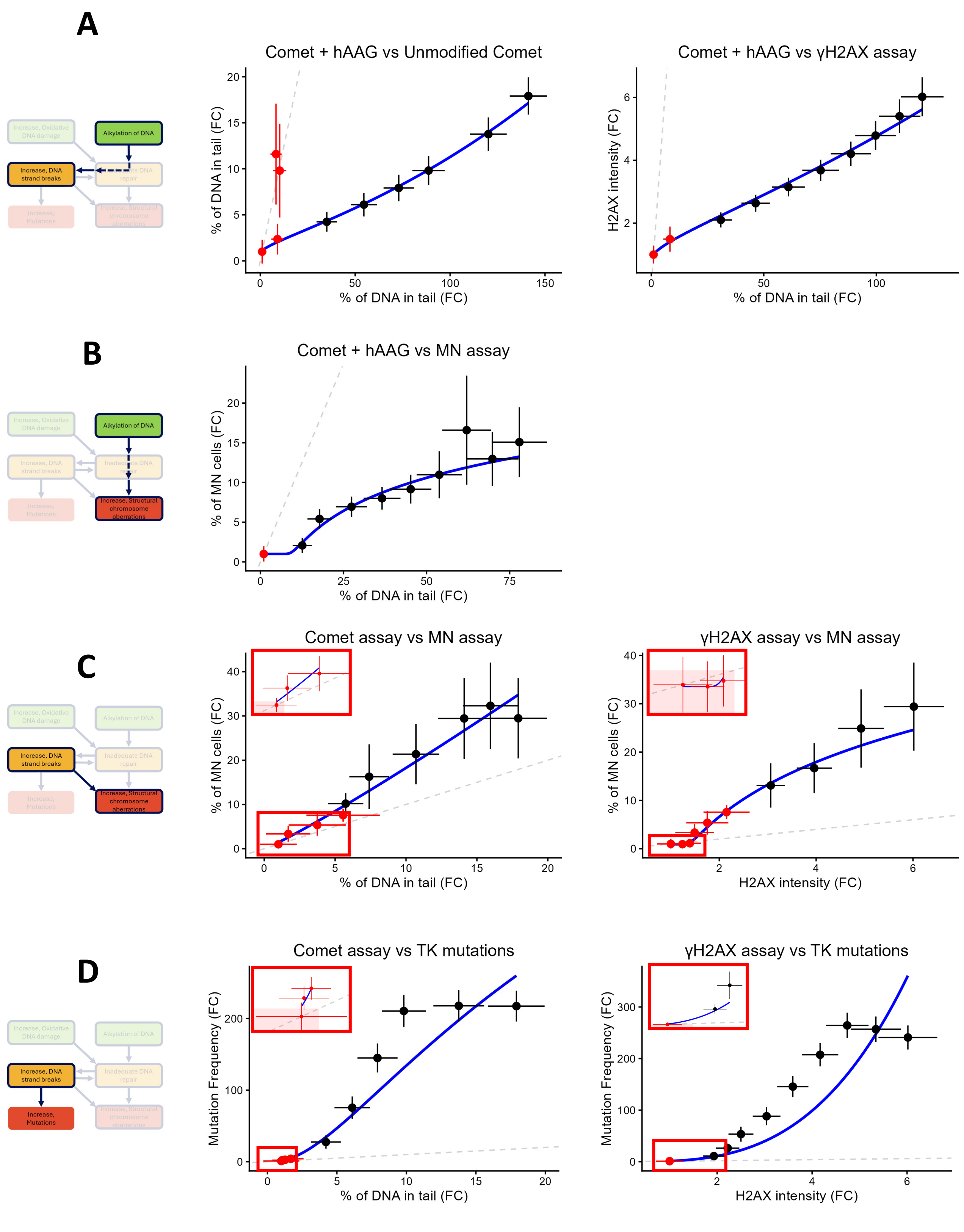
A
Comet + hAAG vs Unmodified Comet
% of DNA in tail (FC)
% of DNA in tail (FC)
Comet + hAAG vs γH2AX assay
H2AX intensity (FC)
% of DNA in tail (FC)
Alkylation of DNA
Increase, DNA strand breaks
B
Comet + hAAG vs MN assay
% of MN cells (FC)
% of DNA in tail (FC)
Alkylation of DNA
Increase, Structural chromosome aberrations
C
Comet assay vs MN assay
% of MN cells (FC)
% of DNA in tail (FC)
γH2AX assay vs MN assay
% of MN cells (FC)
H2AX intensity (FC)
Increase, DNA strand breaks
Increase, Structural chromosome aberrations
D
Comet assay vs TK mutations
Mutation Frequency (FC)
% of DNA in tail (FC)
γH2AX assay vs TK mutations
Mutation Frequency (FC)
H2AX intensity (FC)
Increase, DNA strand breaks
Increase, Mutations

**Figure S5. Response-response modeling for MMS allows for prediction of late stage KEs from upstream KEs.** Fitted response-response models for the following adjacent and non-adjacent KEs: Alkylation of DNA leads to Increase in DNA strand breaks (A), Alkylation of DNA leads to increase in structural chromosome aberrations (B), DNA strand breaks lead to increase in structural chromosome aberrations (C) and DNA strand breaks lead to increase in mutations (D). Responses on all genotoxic endpoints are reported as FCs relative to control. Red dots indicate experimentally matched pairs of mean responses whereas black dots indicate artificial data in the form of means obtained from bootstrap prediction curves through BMC modelling. Error bars indicate the square root of SDs on either observed data (red) or artificial data (black). The leftmost panels indicate the studied KERs, where solid lines are used for adjacent KERs and dashed lines for non-adjacent KERs. In the middle and right panels, the grey dashed line indicates the identity line (y=x). Insets within red boxes show zoom-ins at low responses, with red shaded rectangles denoting the 1.5-fold responses (corresponding with the BMRs).

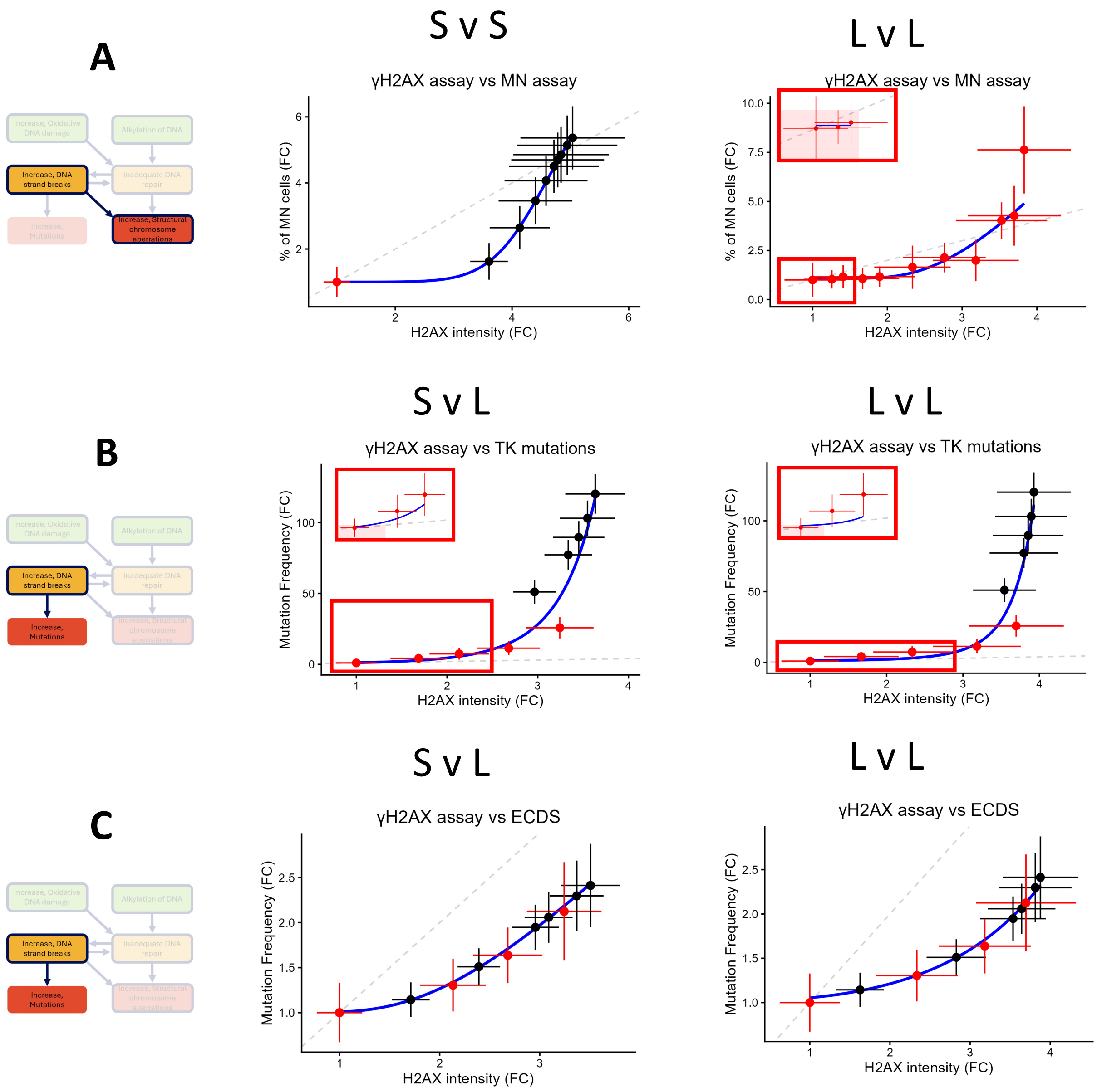

A
B
C
S v S
L v L
S v L
L v L
S v L
L v L
γH2AX assay vs MN assay
γH2AX assay vs MN assay
γH2AX assay vs TK mutations
γH2AX assay vs TK mutations
γH2AX assay vs ECDS
γH2AX assay vs ECDS
% of MN cells (FC)
Mutation Frequency (FC)
H2AX intensity (FC)
Increase, Oxidative DNA damage
Alkylation of DNA
Increase, DNA strand breaks
Inadequate DNA repair
Increase, Mutations
Increase, Structural chromosome aberrations

**Figure S6. Response-response modeling for ENU allows for prediction of late stage KEs from upstream KEs.** Fitted response-response models for the following adjacent and non-adjacent KEs: DNA strand breaks lead to increase in structural chromosome aberrations (A), DNA strand breaks lead to increase in mutations, with mutations measured either through the TK mutation assay (B) or ECDS (C). Titles above the subpanels indicate exposure duration of the underlying datasets, eg: S v L indicates short exposure (≤10 h) vs long exposure (>10 h). Responses on all genotoxic endpoints are reported as FCs relative to control. Red dots indicate experimentally matched pairs of mean responses whereas black dots indicate artificial data in the form of means obtained from bootstrap prediction curves through BMC modelling. Error bars indicate the square root of SDs on either observed data (red) or artificial data (black). The leftmost panels indicate the studied KERs, where solid lines are used for adjacent KERs and dashed lines for non-adjacent KERs. In the middle and right panels, the grey dashed line indicates the identity line (y=x). Insets within red boxes show zoom-ins at low responses, with red shaded rectangles denoting the 1.5-fold responses (corresponding with the BMRs).
